# Replicated Umbilical Cord Blood DNA Methylation Loci Associated with Gestational Age at Birth

**DOI:** 10.1101/749135

**Authors:** Timothy P. York, Colleen Jackson-Cook, Sara Moyer, Roxann Roberson-Nay, Susan K. Murphy, Bernard F. Fuemmeler, Shawn J. Latendresse, Dana M. Lapato, Aaron R. Wolen, Elizabeth K. Do, Catherine Hoyo, Jerome F. Strauss

## Abstract

**Background:** DNA methylation is highly sensitive to *in utero* perturbations and has an established role in both embryonic development and regulation of gene expression. The fetal genetic component has been previously shown to contribute significantly to the timing of birth, yet little is known about the identity and behavior of individual genes.

**Objectives:** The aim of this study was to test the extent genome-wide DNA methylation levels in umbilical cord blood were associated with gestational age at birth (GA). Findings were validated in an independent sample and evidence for the regulation of gene expression was evaluated for *cis* gene relationships in matched specimens.

**Results:** Genome-wide DNA methylation, measured by the Illumina Infinium Human Methylation 450K BeadChip, was associated with GA for 2,372 CpG probes (5% false discovery rate) in both the Pregnancy, Race, Environment, Genes (PREG – Virginia Commonwealth University) and Newborn Epigenetic Study (NEST – Duke University) cohorts. Significant probes mapped to 1,640 characterized genes and an association with nearby gene expression measures obtained by the Affymetrix HG-133A microarray was found for 11 genes. Differentially methylated positions were enriched for actively transcribed and enhancer chromatin states, were predominately located outside of CpG islands, and mapped to genes enriched for inflammation and innate immunity ontologies. In both PREG and NEST, the first principal component derived from these probes explained approximately one-half (58.1% and 47.8%, respectively) of the variation in GA. This assessment provides a strong evidence to support the importance of DNAm change throughout the gestational time period.

**Conclusions:** These results converge on support for the role of variation in DNAm measures as an important genetic regulatory mechanism contributing to inter-individual differences in gestational age at birth. In particular, the pathways described are consistent with the well-known hypothesis of pathogen detection and response by the immune system to elicit premature labor as a consequence of unscheduled inflammation.

## 1 Introduction

Births at less than 37 completed weeks of gestation are preterm and account for most perinatal deaths.^41^ The contribution of both genetic and environmental factors to gestational age at birth (GA) has been established from large twin and family studies evaluating births in Scandinavian^48,87^ and European-American study cohorts.^88,89^ The results of these studies provide point estimates for maternal genetic influences ranging between 13-20% and for fetal genetic sources of 11-35%. While there has been some recent success in the identification of specific additive genetic loci that account for these estimates, the combined influence on risk to preterm birth from these studies has been small.^27,92,93^ Emerging evidence for rare variants that influence risk promises larger effect sizes but for a small subset of births^53,73^ and likely does not contribute directly to observed heritability estimates.^49^

The impact of environmental sources on gestational age at birth is substantial^86^ and varies dramatically across racial groups^12^ that have markedly different preterm birth rates. Only a single twin and family study has directly estimated these influences in African American births and shows that racial differences in preterm birth rates in African Americans versus European Americans can best be explained by environmental, not genetic, factors.^89^ While the impact of non-genetic sources on preterm birth risk is not well understood^11^, further evidence that the variance of GA in African Americans is nearly twice that of European Americans supports the contribution of a heterogeneous array of at least broad categories of environmental exposures.^24^

Whether and how environmental exposures contribute to changes in pathophysiological pathways responsible for triggering early births is an open question. Plasticity in DNA methylation (DNAm) has been observed and numerous environmental risk factors for poor birth outcomes have been reported to be associated with variability in DNAm levels.^39,67^ More directly related to birth outcomes, DNAm changes have been identified in genes relevant to the establishment of pregnancy, the invasion of placental trophoblasts, and fetal development.^8,38,75^ Environmental influence on DNAm levels could initiate from both social exposures and/or pathogenic sources, yet in distinct pathways. For the former, the depolarization of neurons resulting from social environmental signals has been associated with the alterations of the patterns of DNAm and demethylation.^26,50,54^ While the long-term effect of DNAm remodeling due to persistent activation of the HPA-axis and/or the neuronal microenvironment on human development is not clear, it is known that the HPA-axis controls the stress response system that influences the impact of fetal-maternal hormonal signaling on the intrauterine milieu, the maternal-placental interface and fetal physiology.^23,51,72^ For the latter, investigators have recently suggested a provocative case for considering dysregulation in the genetic control of the inflammatory response and innate immune regulation due to pathogenic insults.^65,66,73^

The aim of this study was to test the extent genome-wide DNA methylation levels in umbilical cord blood were associated with GA. The validity of findings were assessed using two independent, racially mixed epidemiological cohorts. The first was the Virginia based Pregnancy, Race, Environment and Genes (PREG) study of samples collected at the Virginia Commonwealth University Health System and the second was from the North Carolina based Newborn Epigenetic Study (NEST) maintained at Duke University. A possible functional relationship between differentially methylated loci with neighboring gene expression was assessed.

## 2 Methods

### 2.1 Samples

#### 2.1.1 Pregnancy, Race, Environment, Genes (PREG) Cohort

The Pregnancy, Race, Environment, Genes (PREG) study was a prospective longitudinal cohort that followed 240 women over the course of pregnancy and has been summarized in *Lapato et. al., 2018*.^39^ All female participants were between 18 and 40 years of age with singleton pregnancies and no diagnosis of diabetes or indication of assisted reproductive technology as verified by medical records abstraction. Both mother and father had to self-identify as either African-American or European-American and be absent of Hispanic or Middle Eastern ancestry. Exclusion criteria at birth included any congenital abnormality, polyhydramnios/oligohydramnios, pre-eclampsia/pregnancy-induced hypertension (PIH)/haemolysis, elevated liver enzymes, low platelet count (HELLP), Rh sensitization, abruptio placentae, placenta previa, cervical cerclage, medically necessitated preterm delivery and drug abuse. Health status was confirmed by medical records abstraction performed by a trained research nurse. At recruitment, before 24 weeks gestational age, women completed a baseline self-report questionnaire to assess demographic characteristics. Women were recruited from Virginia Commonwealth University (VCU) health clinics between 2013-2016. A total of 177 dyads met all criteria and umbilical cord blood specimens were obtained from 135 births. The VCU Institutional Review Board (IRB) approved the study design (HM#14000).

#### 2.1.2 Newborn Epigenetic Study (NEST) Cohort

A second cohort was obtained from a community sample of pregnant women and children who participated in the a North Carolina-based Newborn Epigenetic Study. Pregnant women were recruited from prenatal clinics serving Duke University Hospital and Durham Regional Hospital Obstetrics facilities from 2005 to 2011. Participant identification and enrollment for NEST are described in greater detail elsewhere.^32,46^ Briefly, to be eligible to participate in NEST, participants needed to be at least 18 years or older, pregnant, English and/or Spanish speaking, and intending to use one of two obstetrics facilitates for the index pregnancy to allow for access to labor and birth outcome data. Women completed a self-report or interview-administered questionnaires, which included measures on sociodemographic characteristics, maternal physical health, mental health, and other lifestyle factors. Exclusion criteria included women intending to move before the first birthday of the offspring, relinquish custody of the index child, or who had confirmed human immunodeficiency virus (HIV) infection. Trained personnel abstracted birth information from medical records following delivery, including information on gestational age at birth and other potential covariates. The median GA at enrollment was 12 weeks.

### 2.2 DNA Methylation

#### 2.2.1 PREG DNA methylation

Genomic DNA was isolated from 10 mL whole blood (collected into EDTA tubes) according to standard methods using the Puregene DNA Isolation Kit (Qiagen; Valencia, CA). An aliquot of 1 *µ*g DNA per participant was then sent to HudsonAlpha Institute for Biotechnology for bisulfite conversion (Zymo Research EZ Methylation Kit). Genome-wide methylation was assayed according to the manufacturer’s protocol (Illumina) using the Illumina Infinium Human Methylation 450K BeadChip (Illumina, San Diego, CA, USA), which interrogates 485,764 features. Samples were randomized to arrays and major processing steps to minimize any potential artifactual differences in DNAm patterns that might arise due to batch effects related to processing.

#### 2.2.2 NEST DNA methylation

Ten mL of peripheral blood was collected into EDTA tubes with 1 mL subsequently stored whole. The remainder of the specimen was centrifuged to obtain plasma and the buffy coat layer. Genomic DNA was isolated from the buffy coat using the Qiagen DNeasy Blood and Tissue Kit (Qiagen; Valencia, CA) and then treated with sodium bisulfite using the Zymo EZ DNA Methylation Kit (Zymo Research; Irvine, CA). HumanMethylation450 BeadChip data was generated from the bisulfite modified DNA by the Duke Molecular Center for Human Genetics Shared Resource.

#### 2.2.3 DNAm array data processing

Details of the Illumina Infinium Human BeadChip 450K Array have been previously described^9^ and raw data processing was performed according to best practices reported in recent publications.^57,83^ Intensity values from the scanned arrays were processed using the `minfi` Bioconductor package^2^ in the R programming environment^33,63^. Data processing and analysis was performed separately for the PREG and NEST samples unless otherwise indicated. A summary of sample and probe filtering can be found in Figure 1.

**Figure 1:**
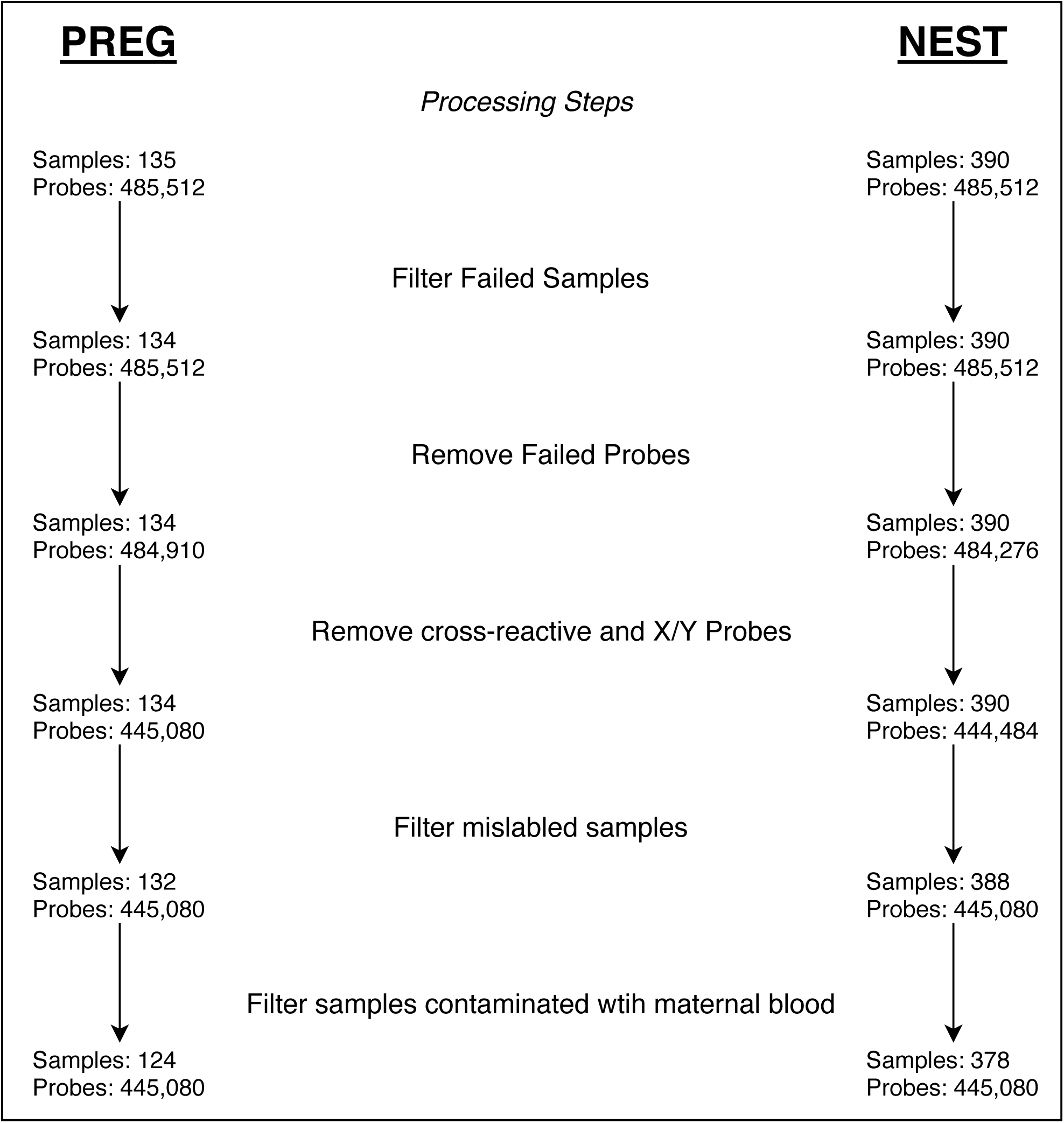
PREG and NEST cohort DNA methylation array probe and sample filtering summary for major processing steps.

Poor quality samples were identified by inspection of sample clustering from a scatter plot of the log median intensities of the raw methylated values against those of the unmethylated values for each array. Good quality arrays tend to group together and poorer quality arrays tend to deviate towards lower median values in both dimensions.^2^ Additionally, beta value density plots from each array were inspected to tag poor performing arrays based on a large deviation from the rest of the samples. Verification of sample identity in the PREG study was made by comparing the correlation of the 65 SNP probes included on the 450k array from fetal samples (umbilical cord blood) with their corresponding maternal samples. Maternal samples were not available for the NEST study but a verification step was performed using similar methods to ensure a unique sample identity. Confirmation of self-reported race was made by sample clustering derived from principal component analysis built upon ancestry informative CpG probes^6^. Maternal blood contamination of umbilical cord blood samples was identified using a panel of CpG markers.^56^

Probes were filtered if they; (1) had a detection P-value of greater than 0.01 in at least 10% of samples; (2) hybridized to either sex chromosome or; (3) have been previously identified as cross-hybridizing.^15^ Probes containing polymorphisms were not removed since the contribution to DNAm variation due to both genetic or environmental sources was of interest as possible etiologic components to the timing of birth. However, additional sensitivity analyses confirmed whether results were enriched for CpG probes containing SNPs. Probe intensities were converted to *β* values which quantifies the proportion each CpG probe is methylated. Quantile normalization adapted to DNAm arrays^77^ was applied to the final set of sample arrays to adjust distributions of type I and II probes on this Illumina platform. This procedure was performed within regions since probe types are confounded across regions and could be expected to have different distributions.^2^

For all statistical tests, the *β* values were transformed using the M-value transformation to promote normality and calculated as a logit transformation of the methylated to unmethylated intensity ratio along with an added constant to offset potentially small values.^17^ Correlations between major experimental factors and the top 10 principal components of M-values across all arrays were inspected to identify extraneous structure that may account for batch effects^45^. ComBat was used to remove average differences across arrays due to slide groupings.^44^ Blood cell proportions were inferred for each sample using the method of Houseman *et. al.* to account for cellular heterogeneity specific to umbilical cord blood.^2,5,31^

#### 2.2.4 DNAm Analytic Model

The analytic aim was to assess the association of GA on DNAm levels derived from umbilical cord blood. This was done in each sample separately following the general form of Equation 1,

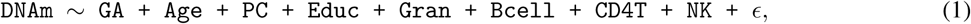

where, `DNAm` is the normalized methyl value, GA is gestational age in days at birth, Age is maternal age in years at enrollment, Educ is maternal education level and *E* was the error term. The cellular component estimates (e.g., `Gran`, `Bcell`, `CD4T`, `NK`, `nRBC`) were included if they correlated with GA in either cohort. Nucleated red blood cell (nRBC) estimates were not included for either cohort consistent with a previously study.^7^ The PC term represented the set of ancestry informative CpG probe set that explain unique variation in GA, which were allowed to differ based on cohort. The PREG sample contained three such terms while NEST included 5, which reflect control for differing population structure between samples. A false discovery rate (FDR) of 5% was used to identify differentially methylated positions (DMP).^82^

#### 2.2.5 DNAm Functional and Regulatory Enrichment

The distribution of significant CpG probes identified to be differentially methylated by GA were examined across functional and regulatory annotations. CpG findings were mapped to known genes^14^ for enrichment of Gene Ontology classifications.^3^ using the `clusterProfiler` package (v3.8.1).^91^ CpG probes were assigned an Entrez identifier if they were within 2 kb upstream or 200 bp downstream of a gene range. Classification functions included biological processes, cellular components, and molecular function, in addition to KEGG pathways. Gene-sets were limited to between 10 and 1,000 members. The R package `GOSemSim` was used to remove redundancy among gene ontology terms due to the directed acyclic nature of the nested GO terms.^90^ Tests for non-random association of CpG probes with CpG island features and ChromHMM chromatin states were based on the AH5086 and AH46969 tracks, respectively, obtained from the AnnotationHub package,^55^ derived from the ENCODE project^29^ CpG island shores were defined as being 2 kb regions flanking CpG islands, while shelves were demarcated as 2 kb upstream or downstream shore regions. For all enrichment evaluations a hypergeometric test for each of these annotations was calculated. When specified, bootstrap methods using 1,000 resamplings were used to estimate 95% confidence intervals. The background set of CpG sites was specific to each cohort as the probes remaining after quality control and filtering (1).

### 2.3 PREG DNAm *cis* Gene Expression

#### 2.3.1 PREG gene expression processing

Total RNA was extracted and the quality evaluated using a previously established sample processing method.^18^ Briefly, RNA was extracted from 10 ml of whole blood and RNA purity was judged by spectrophotometry at 260, 270, and 280 nm. RNA integrity as well as cDNA and cRNA synthesis products were assessed by running 1 *µ*L of every sample in RNA 6000 Nano LabChips on the 2100 Bioanalyzer (Agilent Technologies, Foster City, CA). Biotinylated cRNA was generated with the GeneChip 3’IVT Express kit and 20 *µ*g of the cRNA product were fragmented and 15 mg of the fragmented product were hybridized for 18 to 20 hours into HG-133A microarrays, containing 22,283 probe sets. Each microarray was washed and stained with streptavidin-phycoerythrin and scanned at a 6 mm resolution by the Agilent G2500A Technologies Gene Array scanner (Agilent Technologies, Palo Alto, CA) according to the GeneChip Expression Analysis Technical Manual procedures (Affymetrix, Santa Clara, CA). The overall quality of each array was assessed by monitoring the 30/50 ratios for 2 housekeeping genes (GAPDH and b-actin) and the percentage of ‘present’ genes. The processing of raw data files (Figure 2), including background correction, normalization, and estimation of probe set expression summaries, was performed using the log-scale robust multiarray analysis (RMA) method^34^ as implemented in the affy R package.^21^

**Figure 2:**
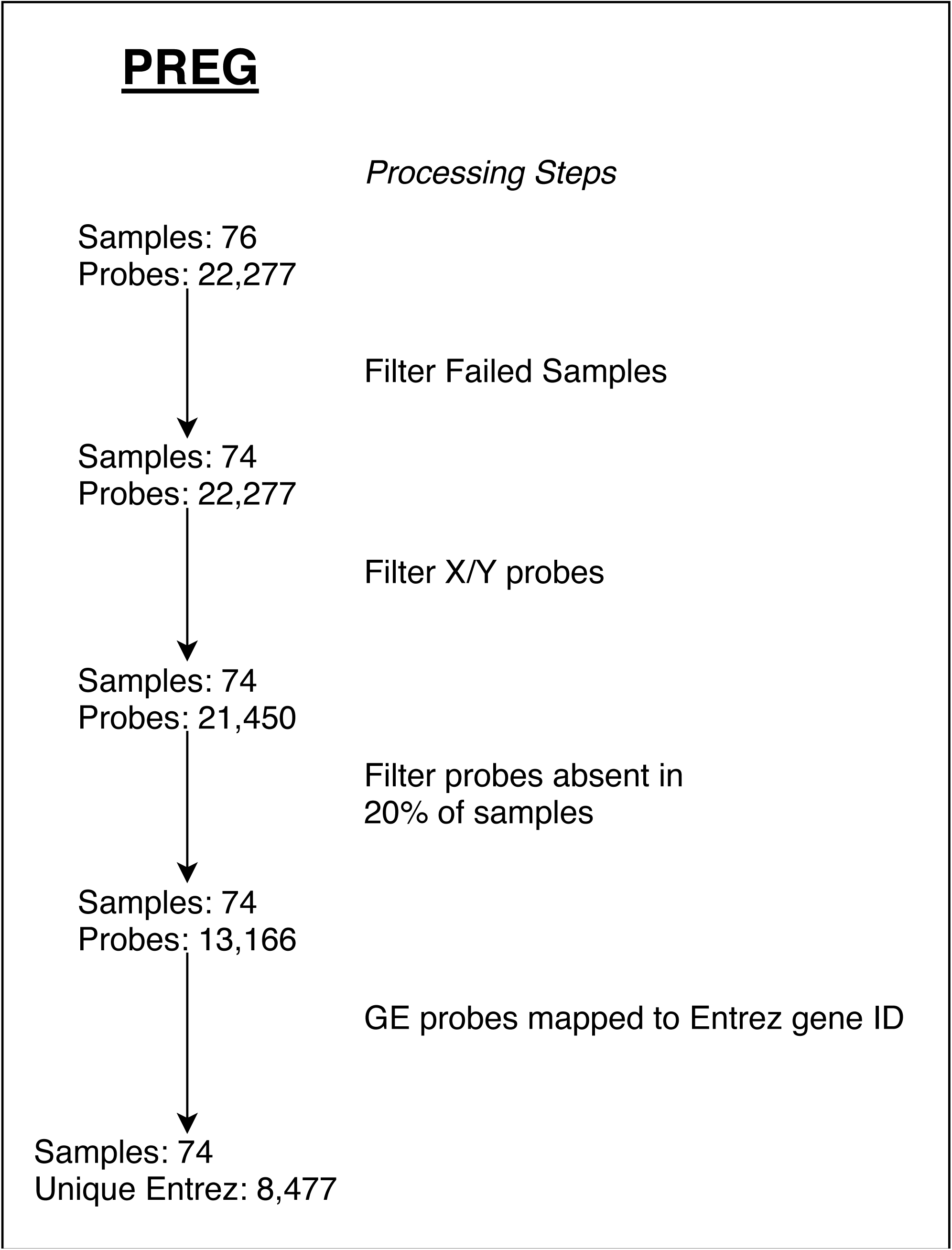
PREG gene expression array probe and sample filtering summary for major processing steps.

#### 2.3.2 *cis* Gene Expression Association

A test of association between gene expression and DMPs characterized to be in *cis* was performed to gain insight into a possible regulatory relationship. DMPs that overlapped between the PREG and NEST samples (Figure 3) were tested against gene expression probes that mapped in *cis*, as:

**Figure 3:**
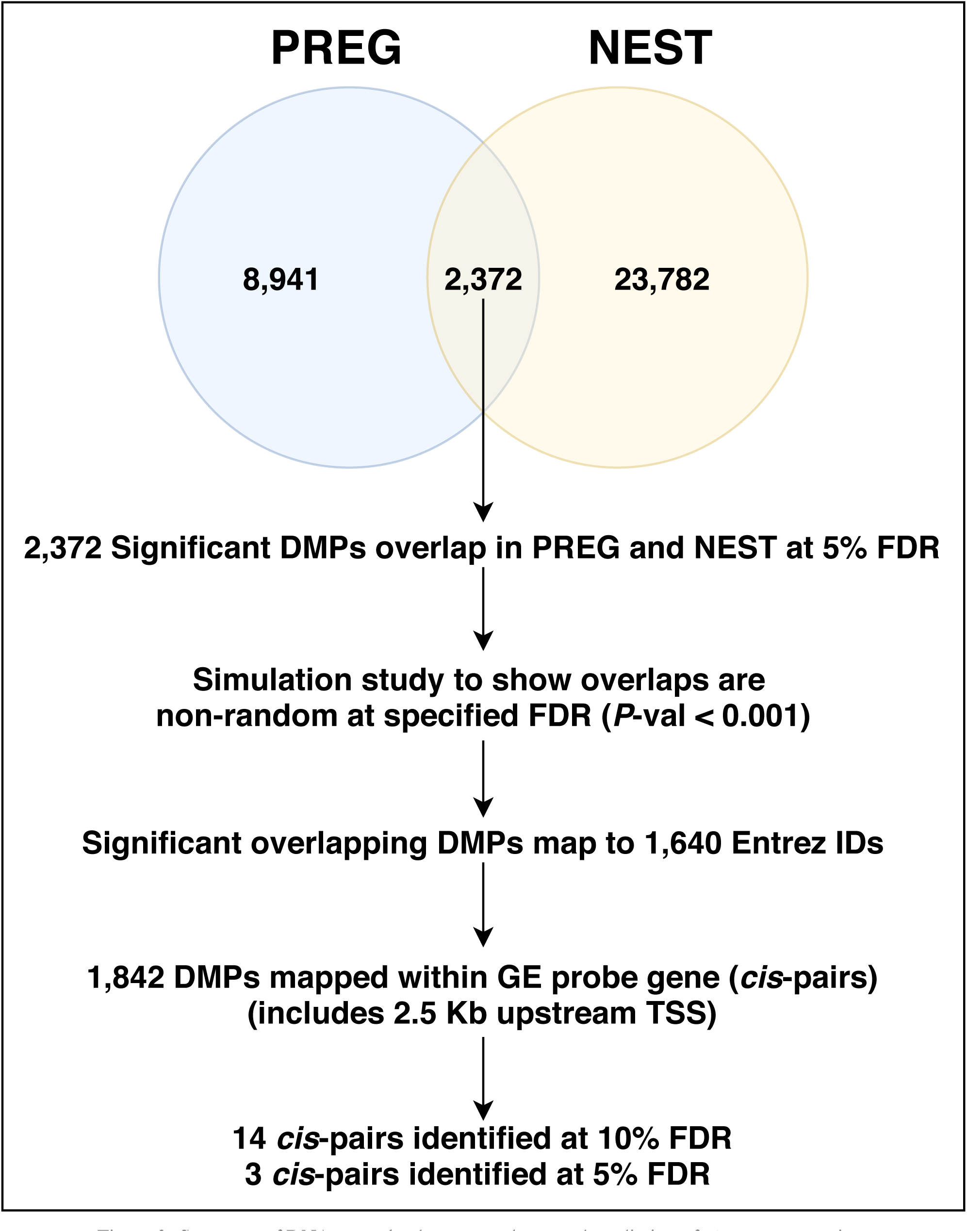
Summary of DNAm overlap between cohorts and prediction of *cis* gene expression.

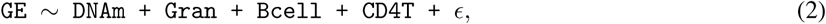

where `GE` was the normalized intensity value of the gene expression probe set, `DNAm` was the normalized intensity of the CpG probe and the remaining were terms for inferred cellular heterogeneity, plus an error term, *ϵ.*

### 2.4 Results

Although not specifically designed as a replication study, the PREG and NEST cohorts share features that allowed for a comparison of certain parameters of our statistical model (Table 1). The PREG participants were born on average about 7 days later than those from NEST (*P*-value < 0.001), which contained a wider range of gestational ages. The PREG cohort was less educated than NEST with fewer participants completing college or receiving a high school diploma or GED (*P*-value < 0.001). Otherwise, characteristics of the sample cohorts did not differ in mean levels of maternal age at birth, self-identified race or sex of the fetus. Gestational age at birth (GA) was not associated with a number of maternal smoking variables in either the PREG or NEST cohort. In PREG neither lifetime smoking (*P*-value = 0.344) nor smoking during pregnancy (*P*-value = 0.359) was associated with GA. The maternal smoking indicator for the NEST cohort also was not associated with GA (*P*-value = 0.093).

**Table 1:**
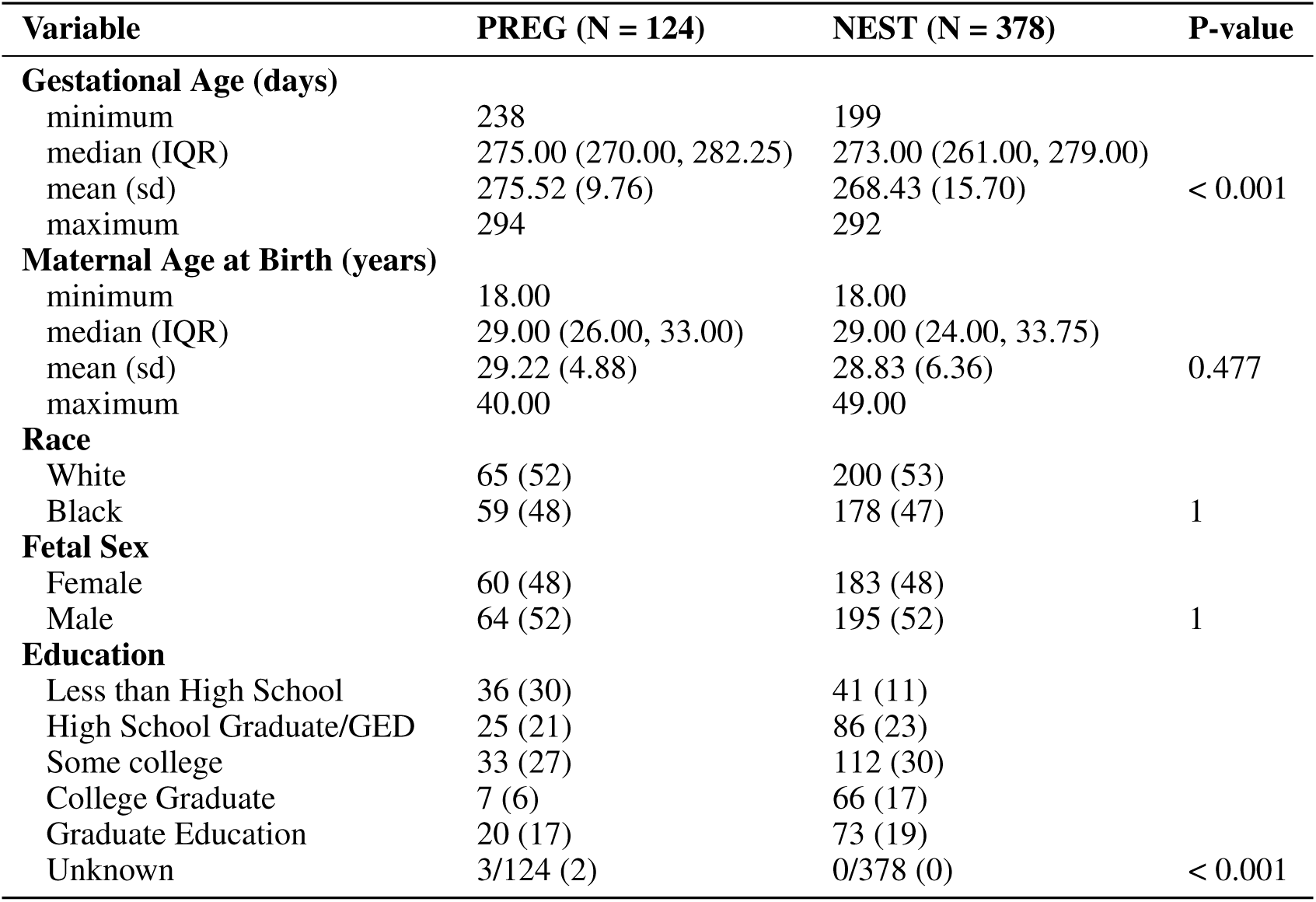
Demographic summary of the PREG and NEST cohorts.

| Variable | PREG (N = 124) | NEST (N = 378) | P-value |
| --- | --- | --- | --- |
| **Gestational Age (days)** |  |  |  |
| minimum | 238 | 199 |  |
| median (IQR) | 275.00 (270.00, 282.25) | 273.00 (261.00, 279.00) |  |
| mean (sd) | 275.52 (9.76) | 268.43 (15.70) | < 0.001 |
| maximum | 294 | 292 |  |
| **Maternal Age at Birth (years)** |  |  |  |
| minimum | 18.00 | 18.00 |  |
| median (IQR) | 29.00 (26.00, 33.00) | 29.00 (24.00, 33.75) |  |
| mean (sd) | 29.22 (4.88) | 28.83 (6.36) | 0.477 |
| maximum | 40.00 | 49.00 |  |
| **Race** |  |  |  |
| White | 65 (52) | 200 (53) |  |
| Black | 59 (48) | 178 (47) | 1 |
| **Fetal Sex** |  |  |  |
| Female | 60 (48) | 183 (48) |  |
| Male | 64 (52) | 195 (52) | 1 |
| **Education** |  |  |  |
| Less than High School | 36 (30) | 41 (11) |  |
| High School Graduate/GED | 25 (21) | 86 (23) |  |
| Some college | 33 (27) | 112 (30) |  |
| College Graduate | 7 (6) | 66 (17) |  |
| Graduate Education | 20 (17) | 73 (19) |  |
| Unknown | 3/124 (2) | 0/378 (0) | < 0.001 |

Initially, the PREG and NEST cohorts contained a total of 135 and 390 umbilical cord blood samples, respectively. Both sets of samples were processed independently following the same general pipeline (Figure 1). A single PREG sample was removed due to low overall median beta intensity values, two samples were removed due to correlation discrepancies with their corresponding maternal samples, and 8 samples were removed due to maternal blood contamination. In NEST, two samples showing high correlations suggesting duplicate samples were removed and 10 samples were removed due to maternal blood contamination. CpG probes were filtered as described in the Methods. Sample selection and probe filtering resulted in 124 samples and 445,080 probes in PREG and 378 samples and 444,484 probes in NEST.

#### 2.4.1 Validation of CpG associations

Univariate tests for the direct association of DNAm with GA were performed as specified by Equation 1. For a 5% false discovery rate (FDR) there were 11,313 CpG probes from the PREG sample meeting this threshold, corresponding to 2.54% of probes tested. At the same FDR, similar tests using the NEST sample resulted in 26,154 CpG probes (5.88% tested). There were 2,372 CpG probes found in both sets that mapped to 1,640 Entrez gene identifiers (Figure 3). Overlapping CpG probes were found to be distributed across the genome, clustered in certain regions, and no strong preference for hyper- or hypo-methylation (Figure 4). Simulation studies were conducted using a combinatorial approach to test whether the observed overlap between samples could be expected by chance. From 10,000 permutations of the data the median number of overlaps expected by chance was 664 CpGs with an interquartile range of 33 confirming the observed degree of overlap was likely driven by the CpGs identified from the GA-DNAm association tested within samples (*P*-value < 0.001).

**Figure 4:**
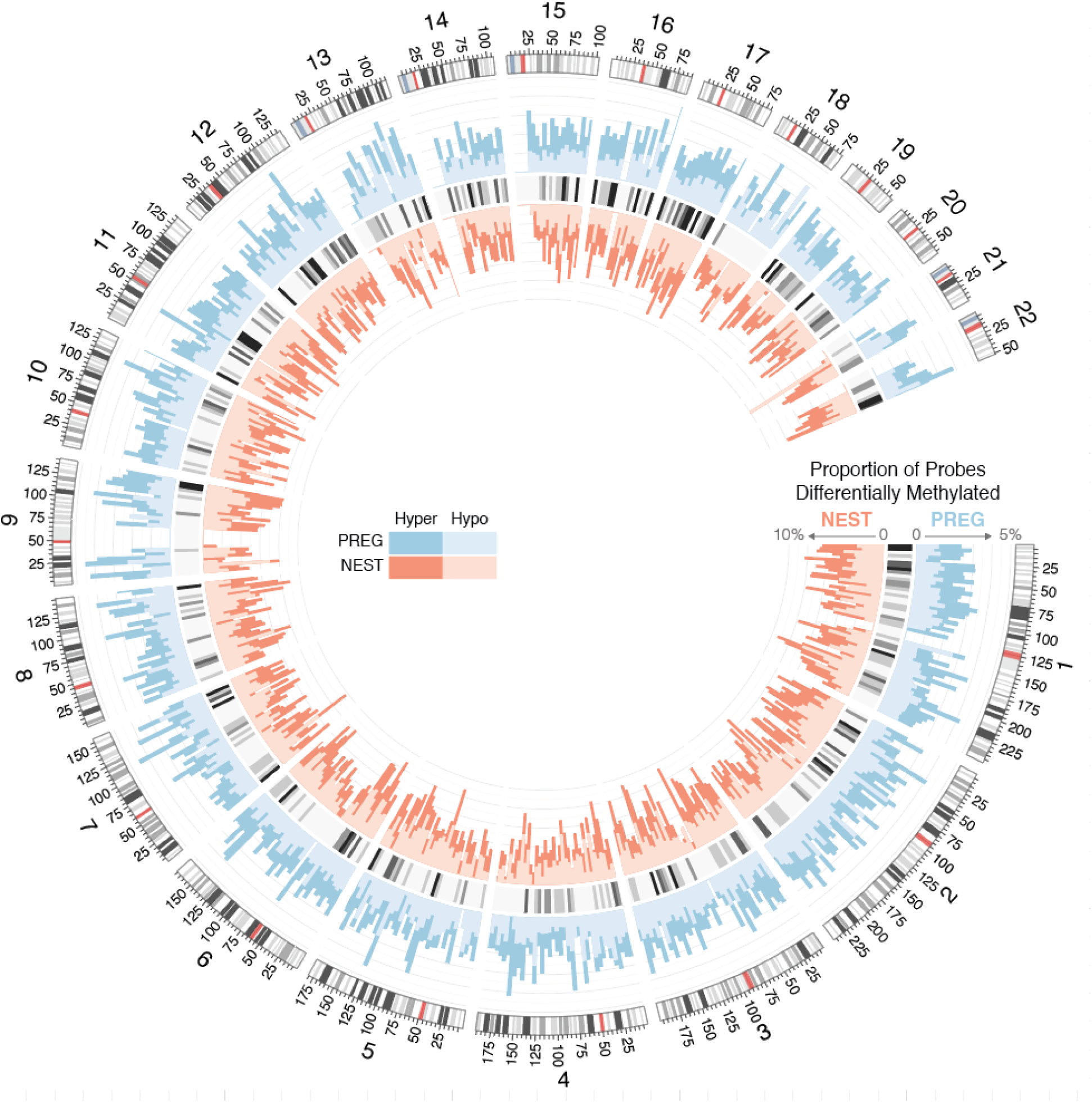
Histograms depict the percentage of CpG probes that are differentially methylated within each cohort across the genome at a resolution of 5 Mb. Hyper- and hypo-methylation is indicated by darker and lighter hues, respectively. The statistical significance of overlapping DMPs within each 5 Mb window was assessed with a one-sided Fisher’s exact test. The FDR adjusted p-values from these tests are visualized by the heatmap separating the PREG and NEST histograms.

Results between samples were found to be highly consistent in terms of direction of association and magnitude of effect. Model coefficients across samples from the 2,372 overlapping probes were correlated at *ρ* = 0.895 (*P*-value < 0.001) and contained the same sign for 97.1% of probes (Figure 5a). A measure of incremental validity was assessed by calculation of variance explained from models with and without the GA term of interest (Equation 1). These *R*-square values were consistent across studies and correlated at *ρ* = 0.390 (*P*-value < 0.001). The mean *R*-square estimated in PREG was slightly higher 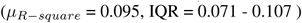 than NEST 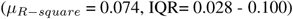; likely driven mostly by sample size differences (Figure 5b).

**Figure 5:**
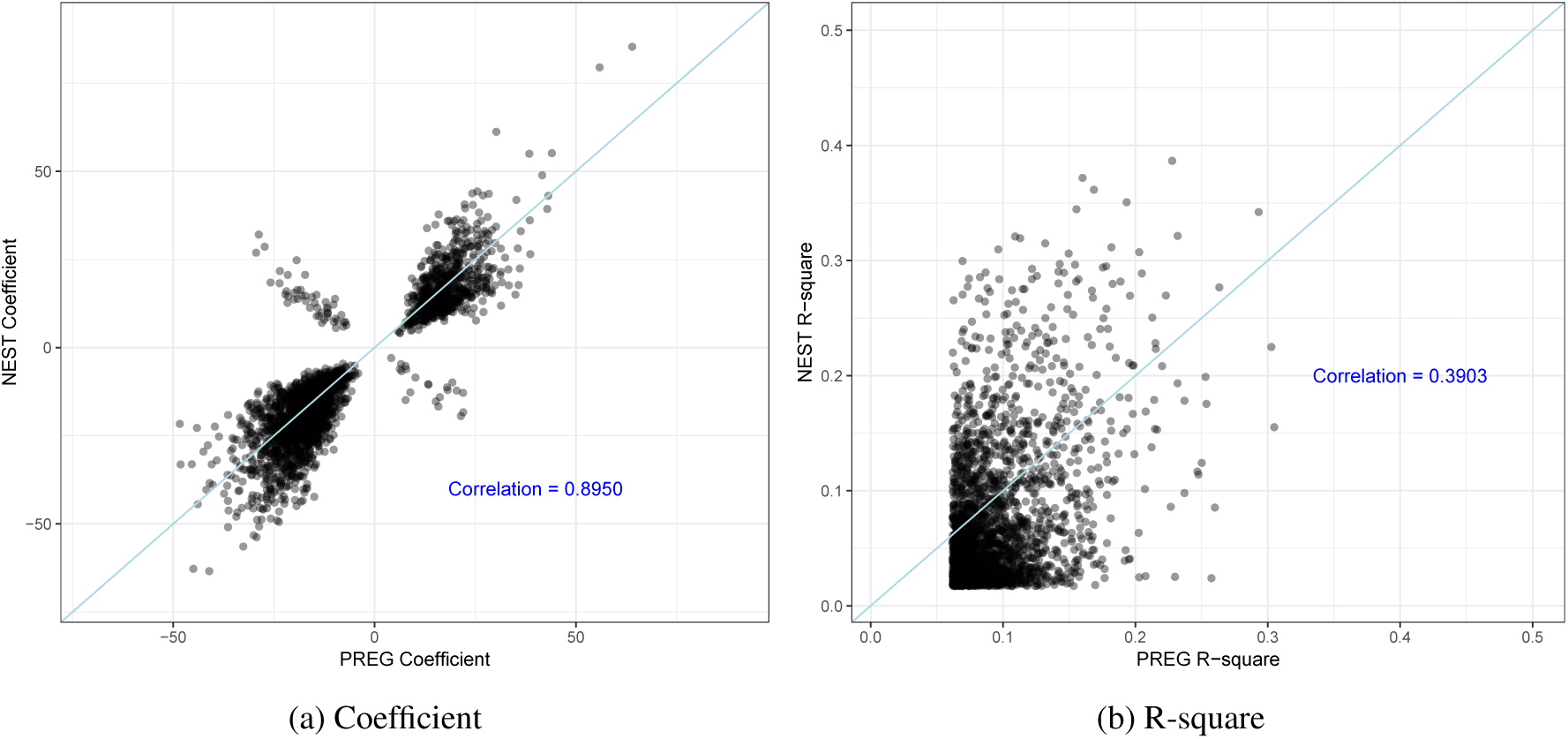
Consistency of model results for overlapping DMPs at an FDR of 5% (N = 2,374) for coefficients (**5a**) and R-square (**5b**) values.

The modest effect sizes observed across a large number of CpG probes could imply redundancy in CpG methylation change either due to the influence of shared pathways or correlated DNA methylation in contiguous CpG sites.^59^ A principal components (PC) analysis was performed to estimate the total amount of variance explained among CpG probes, considering this overlap. For each of the study cohorts, which were analyzed separately, the data matrices of M-values for the overlapping CpG probes were provided as input after adjusting for covariates as specified similarly in Equation 1. The first PC in each sample accounted for 15.5% and 18.9% of the total DMP variance in PREG and NEST, respectively. GA was then regressed on the first derived PC for each sample separately and resulted in a consistent *R*-square value across samples of 0.581 in PREG and 0.478 in NEST.

#### 2.4.2 Enrichment and biological relevance

The 2,372 CpGs that overlapped in each sample at a 5% FDR mapped to 1,640 unique genes and associated promoter regions. This consisted of 1,227 CpGs within gene ranges and 704 falling in promoter regions. Gene-based enrichment was conducted to provide an overview of DNAm contributions at the level of function (*Molecular Function*), where gene products are active (*Cellular Component*), pathways of multiple gene products (*Biological Processes*) and curated pathways (*KEGG*).^91^ The results of enrichment tests yielded 21 significant categories filtered at a FDR of 1% (Figure 6). These generally consisted of gene ontology groups enriched for inflammatory activation and immune response ranging from approximately 1 to 6% of DMPs mapped to genes. Along this same functional theme, KEGG pathways were enriched for Th17 cell differentiation.

**Figure 6:**
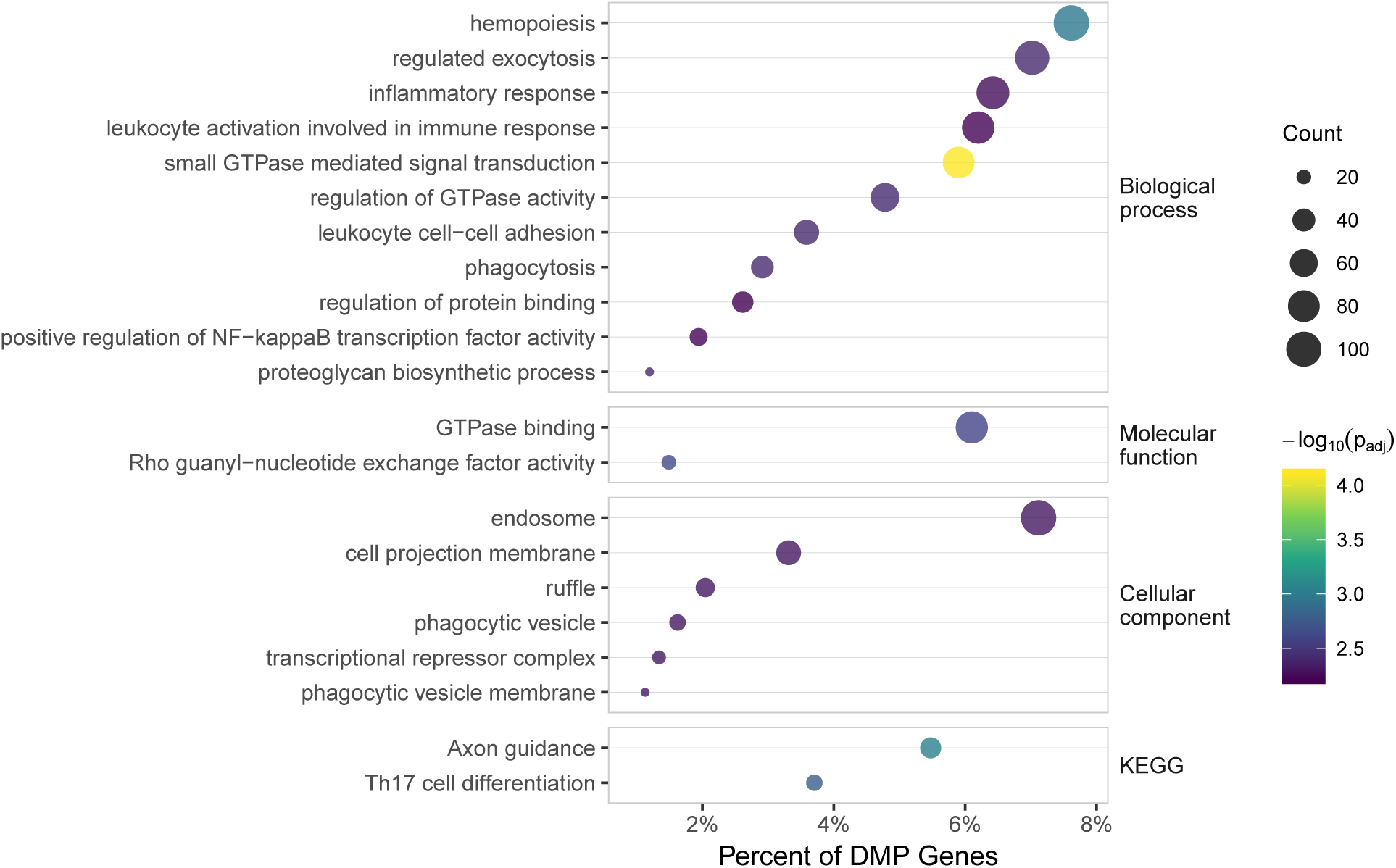
Summary of gene-based enrichment for Gene Ontology groups and KEGG pathways.

The set of overlapping CpG probes was found to have a nonrandom pattern of association with CpG island and chromatin based annotations. CpG island enrichment was seen for both north (*P*-value = 0.002) and south (*P*-value = 0.001) shore regions while depleted in CpG islands (*P*-value = 0.001) (Supplemental Figure 7). ChromHMM category enrichment (Supplemental Figure 8) included Flanking Active TSS (*P*-value = 0.001), Transcription at gene 5’/3’ (*P*-value = 0.001), Weak Transcription (*P*-value = 0.007), Genic Enhancers (*P*-value = 0.001) and Enhancers (*P*-value = 0.001). Significant ChromHMM depletion was observed in Active TSS (*P*-value = 0.001), Strong Transcription (*P*-value < 0.003), Heterochromatin (*P*-value = 0.001), Bivalent/Poised TSS (*P*-value = 0.006), Flanking Bivalent TSS/Enh (*P*-value = 0.040), Quiescent/Low (*P*-value = 0.001).

**Figure 7:**
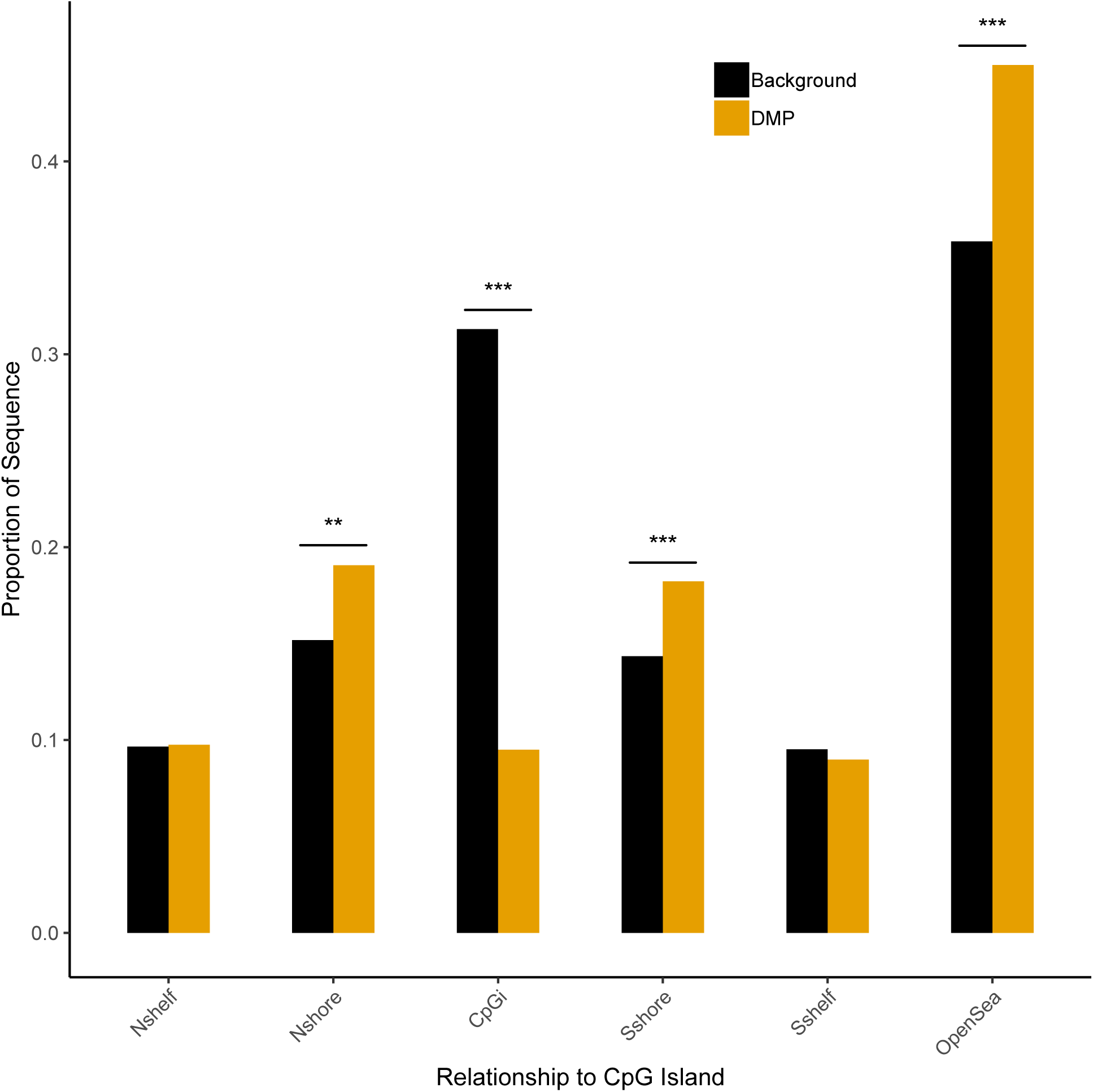
Localization of the differentially methylated positions to CpG island characteristics. The Background (black) denotes the union of CpG probes tested in both cohorts. DMP (gold) was the intersection of overlapping DMPs identified in both cohorts (N = 2,374). * = *P*-value < 0.05; ** = *P*-value < 0.01; *** = *P*-value < 0.001

**Figure 8:**
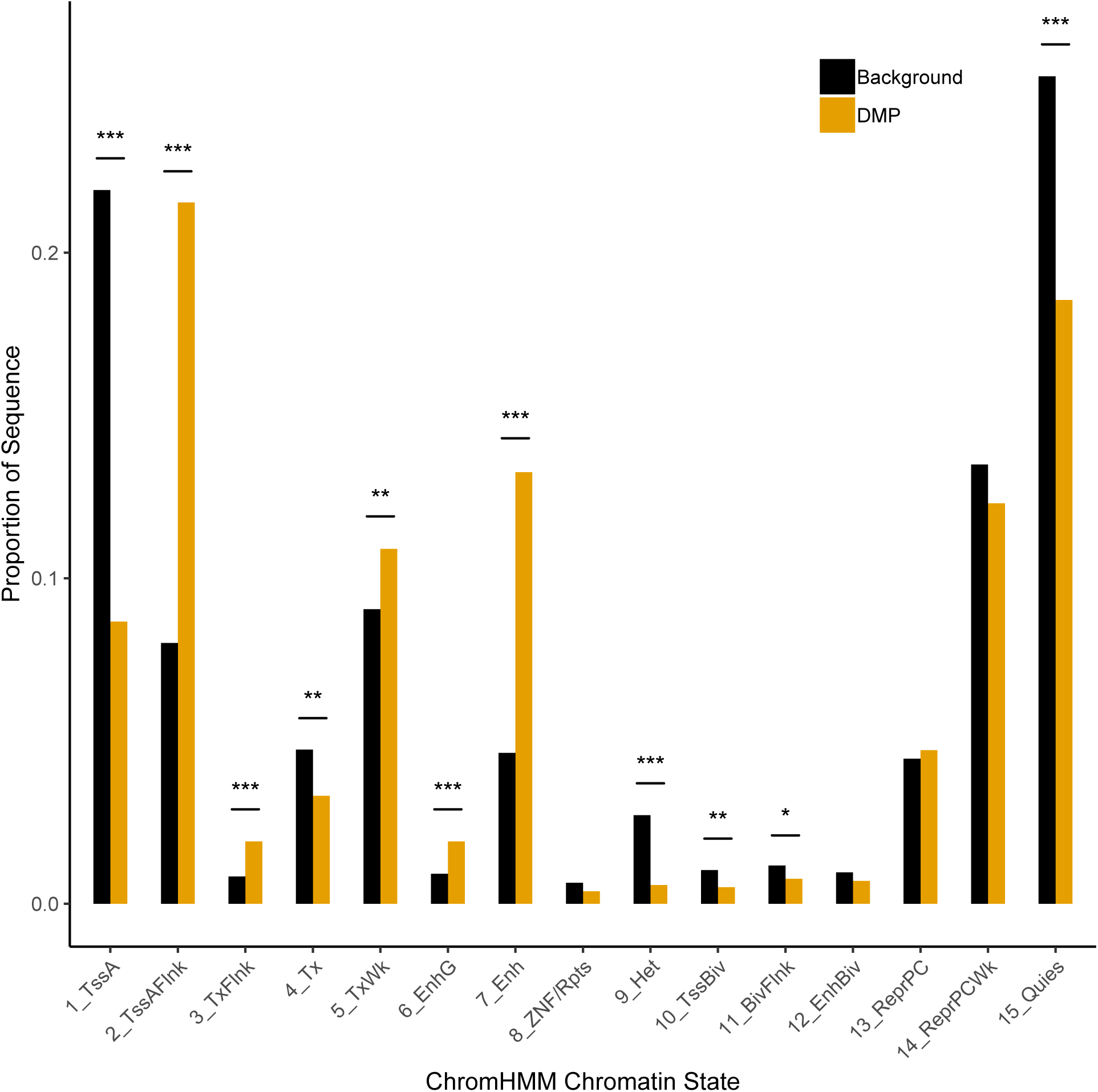
The proportion of DNA sequence, measured by CpG probes, by ChromHMM defined chromatin characteristics. The Background (black) denotes the union of CpG probes tested in both samples. DMP (gold) was the intersection of overlapping DMPs identified in both samples (N = 2,374). The 15 chromatin states are numbered and abbreviated as: 1_Active Transciption Start Site (TSS), 2_Flanking active TSS, 3_Transcription at gene 5’ and 3’, 4_Strong transcription, 5_Weak Transcription, 6_Genic enhancer, 7_Enhancer, 8_ZNF genes and repeats, 9_Heterochromatin, 10_Bivalent/Poised TSS, 11_Flanking bivalent TSS/Enh, 12_Bivalent enhancer, 13_Repressed PolyComb, 14_Weak repressed PolyComb, and 15_Quiescent/low. * = *P*-value < 0.05; ** = *P*-value < 0.01; *** = *P*-value < 0.001

#### 2.4.3 DMP Association with Gene Expression

Of the 124 PREG specimens retained for DNAm analysis, only 76 also had gene expression array data available. From the set of 76 gene expression arrays, two specimens were removed due to poor sample quality. and 13,166 probesets were present in at least 20% of samples, leaving probesets that mapped on to 8,477 unique Entrez gene identifiers (Figure 2). The 2,372 significant overlapping DMPs mapped to 1,842 genes (including 2.5 Kb upstream from the TSS) in *cis*, which also contained gene expression probesets. Statistical tests were performed on the gene level regardless of the genomic location of the CpG methylation probe or gene expression probesets (Equation 2). This set of association tests resulted in 14 *cis*-pairs at a FDR of 10% (Table 2). The median *R*-square for associations was 0.150 (range = 0.106-0.395) and approximately one-half (8 of 14) of the associations showed an inverse relationship between gene expression and DNAm values (higher expression with lower methylation values). Only the *EXT1* gene contained multiple mappings of CpG probes and gene expression probesets.

**Table 2:** DMP *cis* Gene Expression Association.

| Gene | Entrez ID | Affy Probeset | CpG Probe | Chrom. | CpG Position | Coef | P-value | R-square<sup>1</sup> | Gene Relevance |
| --- | --- | --- | --- | --- | --- | --- | --- | --- | --- |
| TRIM33 | 51592 | 212435_at | cg26410133 | chr1 | 115051834 | -2.309 | 0.001 | 0.156 | Regulates proinflammatory function of Th17 cells<sup>74</sup> |
| CD247 | 919 | 210031_at | cg15518113 | chr1 | 167400121 | -1.124 | 0.000 | 0.395 | Impaired immune response<sup>35</sup> |
| PLCH1 | 23007 | 214745_at | cg26690511 | chr3 | 155422103 | 0.283 | 0.001 | 0.145 | Cord blood association with PTB<sup>20</sup> |
| F2RL1 | 2150 | 213506_at | cg00499700 | chr5 | 76116088 | -1.852 | 0.001 | 0.151 | Regulation of myometrial function at labor and preterm labor<sup>58</sup> |
| KEL | 3792 | 206077_at | cg17784922 | chr7 | 142659425 | -1.242 | 0.001 | 0.113 | Anti-Kell associated with increased risk for PTB<sup>19</sup> |
| EXT1 | 2131 | 201995_at | cg20547777 | chr8 | 119086580 | 0.841 | 0.000 | 0.141 | Necessary for heparan sulfate elongation which enhances neutrophil infiltration<sup>13</sup> |
| EXT1 | 2131 | 214985_at | cg20547777 | chr8 | 119086580 | 0.669 | 0.000 | 0.205 | " |
| EXT1 | 2131 | 201995_at | cg16009311 | chr8 | 119086762 | 0.876 | 0.000 | 0.149 | " |
| EXT1 | 2131 | 214985_at | cg16009311 | chr8 | 119086762 | 0.668 | 0.000 | 0.198 | " |
| SIGIRR | 59307 | 218921_at | cg08869273 | chr11 | 413594 | -1.185 | 0.001 | 0.136 | Negative regulation of TLR and IL-1 receptor signalling to illicit immune response<sup>62</sup> |
| CHD4 | 1108 | 201184_s_at | cg20600850 | chr12 | 6712515 | 0.889 | 0.000 | 0.148 | Nucleosome remodeling and deacetylase complex<sup>47</sup> |
| ART4 | 420 | 207220_at | cg20967028 | chr12 | 14996272 | -1.696 | 0.000 | 0.106 | ADP-ribosyltransferase 4 (Dombrock blood group) |
| CTDP1 | 9150 | 205035_at | cg24998981 | chr18 | 77441810 | -0.878 | 0.000 | 0.175 | CTD phosphatase subunit 1 |
| CST7 | 8530 | 210140_at | cg19204859 | chr20 | 24933798 | -0.875 | 0.000 | 0.159 | Innate immunity by affecting cytotoxicity of NK cells<sup>61</sup> |

<sup>1</sup>R-square is the incremental validity of the CpG predictor.

### 2.5 Discussion

The complexity of data from an epigenome-wide association study (EWAS) precludes a direct assessment of CpG sites that may be etiologically involved in a biological process. This is in contrast to methods that for the most part seek only to identify predictors of disease outcome regardless of etiologic contribution.^22,36,42^ The potential to detect a CpG association signal that exceed expectations for a typical genome-wide association study (GWAS)^9^ should be counterbalanced by the dynamic nature of DNAm whose site-specific measurements can be influenced both by biological variability and random measurement error.^52^ Even small variations in methodological approach can lead to large changes in feature selection across the thousands of CpG probes interrogated. Recommendations made for the conduct of EWAS stress the generation and assessment of multiple criteria to build confidence in identification of robust results.^25,40,52,76^ The approach of the present study focused on the identification of replicated CpG sites associated with GA that were reproduced in two independent studies, thereby building evidence for a potential transcriptional regulatory function.

#### 2.5.1 Replication of Differentially Methylated Positions

The relationship between GA at birth and DNAm was characterized primarily based on overlapping results from two epidemiological cohorts comprised of pregnant women and their infants. There may be some question as to whether this represents a replication or validation of results in the sense that a full EWAS was performed in both samples instead of confirming individual results. Furthermore, verification of results using a different technique was not performed. Yet, one of the strengths of this study were that both cohorts were collected specifically for birth outcomes research (i.e., not convenience samples). Also, DNA was collected and processed by independent research groups thus eliminating laboratory technical bias that could influence both samples, including sources of major batch effects known to confound statistical analysis.^45^ In order to identify potential significant DMPs in a replication study, the use of FDR would not be appropriate since only a restricted range of effect sizes would be used to estimate the replication result FDR.^78^ Other multiple test correction techniques, including Bonferroni, would likely be too conservative for a large number of results for the same reasons for selecting the FDR in the first place. One strategy would be to verify results using an alternative technology of higher resolution, such as sequencing bisulfite converted DNA, for more in depth studies of a specific chromatin region. For instance, future studies verifying a subset of these results in placental trophoblast cells would provide strong evidence that biologically important changes were detected. The current strategy essentially repeats the EWAS in an independently collected sample and a unified analytic approach applied to both sets of data allows for a straightforward comparison of results.

The consistency of DMP findings across the two cohorts was assessed in multiple ways. The degree of overlap in samples was found to be non-random by simulation study, concordant in direction of change, and similar in size of effects. The NEST study analysis yielded approximately twice the number of significant DMPs at a FDR of 5%. This difference was most likely due to the increased statistical power of the larger NEST sample (about 3 times larger) and can be seen by the wider range of effect sizes detected compared to PREG DMP results (see Figure 5b). An assessment of incremental validity revealed a consistent and moderately sized influence across DMPs for both cohorts. The mean *R*-square estimates of 9.5% and 7.4% suggest a biologically significant amount of variance explained by most DMPs. Regardless, there should be some appreciation that the DMPs identified could exert their influence through redundant biological pathways or more simply be correlated across participants. The derivation of principal components across DMPs and their relationship to gestational age provided a global summary of DNAm influence. In both PREG and NEST the first PC explained approximately one-half (58.1% and 47.8%, respectively) of the variation in GA. This assessment provides a strong justification for the importance of DNAm change throughout the gestational time period.

#### 2.5.2 Biological Relevance

The results of several studies document the relationship between DNAm and chronological age,^1,59^ as well as measures for the biological deviation from age.^28,30^ The first 40 weeks of this distribution has, of course, been targeted as a critical period in establishment of the utero-placental interface, embryonic development and transmission of signal to initiate labor onset. Investigators have observed a direct association of GA on newborn/umbilical cord DNAm^43,60,68,71,84^, including supervised algorithms designed to predict GA.^10,36,42^ These studies are consistent with integrative approaches that have shown involvement of inflammatory and immune-related pathways^4,22,37^, and the larger picture of these pathways as being integral to the onset of labor.^65,66,73^ Of the many potential antecedents, the literature defines a strong pathophysiological link for the causal role of inflammation on preterm birth.^64,66,80^ While most histopathological cases of inflammation (e.g., chorioamnionitis) are sub-clinical, detection by molecular signatures responsible for the activation and enhancement of the innate immune response may provide an early warning sign.^66^

Findings from this study also support the inflammatory activation of the immune system. Not only were genes with mapped DMPs enriched for these ontological categories, but a select number of sites were found to have evidence of a gene regulatory role. Although all replicated DMP findings could potentially be informative, those found to have a direct association with *cis* gene expression values could be considered to have stronger empirical support. Of these 11 unique genes, there were 8 (*TRIM33, CD247, KEL, EXT1, SIGIRR, CST7, F2RL1, ART4*) with previous evidence of an inflammatory/immune function. For example, *TRIM33* is expressed in the placenta and has been shown to mediate the inflammatory function of Th17 cells by inducing *IL-17* and suppressing *IL-10*.^74^ The IL-10 protein has been previously described as a pleiotropic regulator of preterm birth critical in balancing the anti- and pro-inflammatory response at the maternal-fetal interface.^16^ The maintenance of this inflammatory response is also influenced by the placentally expressed *TRIM33* that suppresses IL-10, which in turn promotes the proinflammatory function of Th17 cells.^74^ *SIGIRR* is a negative regulator of innate immune response via the Toll-like receptor/IL-1R signaling pathway.^69,79^ Finally, *EXT1* is a heparan sulfate copolymerase which enhances neutrophil response to IL-8.^81^ IL-8 is constitutively produced by the placenta and enhanced at the fetomaternal interface characterized by neutrophil infiltration in cases of chorioaminionitis.^70^ A broad interpretation of these results converge on reasonable support for the role of variation in DNAm measures as an important genetic regulation mechanism contributing to inter-individual differences in GA. In particular, the pathways described are consistent with the well-known hypothesis of pathogen detection and response by the immune system to elicit premature labor as a consequence of unscheduled inflammation.

DNAm by itself is an imperfect measure of gene activity and does not imply direction of causation in cohort designs. Few integrative studies have been attempted for quantitative traits and little is known about the complex relationship between DNAm and gene expression on a genome-wide scale.^85^ Yet, the integration with other platforms can support hypotheses of causal inference and aid in interpretation, especially when matched to the same samples and time points. The present study could only test for these relationships within the PREG sample which was presumably under-powered for a genome-wide test of all possible *cis* relationships. For similar reasons, the set of all possible *trans* tests was not attempted (i.e., association of DMP sites with any gene expression probe greater than 2 Kb from TSS), which is the likely mechanism of regulation for transcription factors. The approach taken in this study provides a strong *prima facie* test case for integration studies across genomic platforms and, to the best of our knowledge, represents the first such example in studies of birth outcomes. The DMP relationships reported here, validated and mapped alongside gene expression variation, represent a promising direction forward for EWAS studies that have matured past the point of merely descriptive reports. An exciting avenue for future studies would be the development of more extensive models that include measures of environmental exposures (more specifically, preterm birth risk factors) to test hypotheses of their influence on gene expression regulation through DNAm changes.^39^

### 2.6 Conclusion

Current literature supports a “complex, multifactorial framework” for the initiation of labor.^73^ DNAm provides an anchor point for biomedical research directed towards the integration of input streams originating from both genetic and environmental sources. This includes consideration for DNAm plasticity in the presence of environmental exposures along with the moderation of locus-specific DNAm by allelic differences (i.e., *mQTL*). Genome-wide DNAm array studies have been shown to replicate and demonstrate appreciable effect sizes detectable from moderately sized cohorts. The results presented in this study show a convergence of significant DNAm findings from two independent, epidemiological cohorts, along with evidence of DNAm association with neighboring gene expression enriched for hallmark features of labor onset. Due to the tissue specific nature of transcriptional control, including transcription factors, future studies should aim to confirm these findings in a tissue more directly related to parturition.

## 3 Declarations

### 3.1 Acknowledgements

The Pregnancy, Race, Environment, Genes (PREG) longitudinal study was supported by the NIHMD (P60MD002256, PI: York, Strauss). The NEST study was funded by the NIEHS (R21ES014947, PI: Hoyo) and NIEHS (R01ES016772, PI: Hoyo) and the NIDDK (R01DK085173, PI: Hoyo). The use of REDCap was supported by Clinical and Translational Science Award (CTSA) award No. UL1TR000058 from the National Center for Advancing Translational Sciences.

### 3.2 Data sharing

Data sharing is limited by Institutional Review Board (IRB) agreements and participant consent forms, which restrict openly sharing individual-level DNAm measures. Individuals interested in data access or collaboration are encouraged to contact Dr. Timothy P. York

### 3.3 Author contributions

TPY planned and carried out the analysis and wrote the initial manuscript draft. TPY and JFS conceived of and secured funding for the Pregnancy, Race, Environment, Genes (PREG) study. SJL, RRN and DML consulted on the design of statistical models. AW performed the bioinformatic analyses. The PREG specimen processing was overseen by CJC. SM initiated the collection of fetal cord blood samples in PREG. BFF and ED initiated the NEST validation study. SMK and CH provided consultation on the Newborn Epigenetic Study (NEST). All co-authors reviewed the manuscript and approved the final version.

### 3.4 Conflicts of interest

The authors report that they have no conflicts of interest.

### 3.5 Abbreviations

DNAm: DNA methylation
PREG: Pregnancy, Race, Environment, Genes Study
NEST: Newborn Epigenetic Study
TSS: Transcription start site
DMP: Differentially methylated position
EWAS: Epigenome-wide assocation study
GWAS: Genome-wide association study
CpG: Cytosine-Guanine dinucleotide
SNP: Single nucleotide polymorphism
mQTL: Methylation quantitative trait loci
ChromHMM: Chomatin hidden Markov model

